# Intracorneal delivery of HSV-targeting CRISPR/Cas9 mRNA prevents herpetic stromal keratitis

**DOI:** 10.1101/2020.02.08.934125

**Authors:** Di Yin, Sikai Ling, Dawei Wang, Dai Yao, Hao Jiang, Soren Riis Paludan, Jiaxu Hong, Yujia Cai

## Abstract

Herpes simplex virus type 1 (HSV-1) is a leading cause of infectious blindness. Current treatments for HSV-1 do not eliminate the virus and are incapable of modulating the virus reservoir. Here, we target HSV-1 genome directly using mRNA-carrying lentiviral particle (mLP) that simultaneously delivers spCas9 mRNA and two viral genes-targeting gRNAs (designated HSV-1-erasing lentiviral particles, HELP). We showed HELP efficiently blocked HSV-1 replication in both acute and recurrent infection models, and prevented occurrence of herpetic stromal keratitis (HSK). We further showed retrograde transportation of HELP from corneas to trigeminal ganglia (TG) where HSV-1 established latency and found evidence of HELP modulating herpes reservoir. Additionally, the potent antiviral activity of HELP was also replicable in human-derived corneas. These results strongly support clinical development of HELP as a new antiviral therapy and may accelerate mRNA-based CRISPR therapeutics.

---

Herpes simplex virus type I (HSV-1) is among the most common human viruses with 50-90% of the world population being sero-positive^1^. It belongs to alpha-subfamily of herpesviruses, which are enveloped viruses carrying double-stranded DNA^2^. HSV-1 infection can cause a wide variety of diseases including herpes simplex encephalitis, which has a high mortality if untreated^3^. HSV-1 infection in the cornea can cause herpetic stromal keratitis (HSK), which is the leading factor for infectious blindness^4^. After primary infection and production replication in corneal epithelium, HSV-1 travel through ophthalmic nerves in retrograde direction to trigeminal ganglions (TG) where they establish a latent reservoir^4^. Under certain stimuli, the latent viruses in the TG can re-activate, leading to the recurrence and aggravation of the disease. Globally, it is estimated that 1.5 million episodes of ocular HSV each year and 40,000 people get visual disability^4^.

The first-line treatment option for HSV-1 infection is acyclovir (ACV) and its analogs, all targeting to the viral DNA polymerase^5^. The effectiveness of these drugs is frequently inadequate and they suffered from (i) frequent drug resistance including emergence of mutants^6-8^, (ii) ineffective for stromal and endothelial infection^4,9,10^, (iii) side effects including nephrotoxicity induced by prolonged use^11-14^. For better alternatives, antibodies, peptides and small molecular inhibitors are under development^15^. However, none of these strategies are able to disrupt the virus or modulate its reservoir– therefore, incapable of preventing recurrence.

CRISPR cleaves on genomes directly and has shown therapeutic potential in a variety of disease models, in most cases genetic diseases^16-21^. Its therapeutic potential on infectious diseases is promising, but still in its infancy due to lack of strong safety and efficacy evidence supporting for clinical application^22^. So far, no existing studies have shown the therapeutic efficacy of CRISPR *in vivo* against HSK^23^.

In this study, we designed a gRNA expression cassette targeting to two essential genes of HSV-1 -UL8 and UL29, simultaneously, and co-packaged with spCas9 mRNA to mRNA-carrying lentiviral particle (mLP) (Fig. 1a, 1b and 1c). As the resulting lentiviral particle was designed to destroy the genome of HSV-1, we therefore designate it HSV-1-erasing lentiviral particles (HELP). The gRNA expression cassette is reverse transcribed and maintained as circular episomal DNA as integration-defective lentiviral vector (IDLV) (Fig. 1b). As the UL8 gRNA is cloned into the ΔU3 region of long terminal region (LTR), it will be copied from 3’-LTR to 5’-LTR during reverse transcription (Fig. 1b). We produced the HELP by co-transfection of 6 plasmids to 293T cells and harvested the particles by ultracentrifugation (Fig. 1c). As controls, we also produced mLP expressing single gRNA-either UL8, UL29 or scramble (non-targeting gRNA). To verify whether HELP was indeed capable of inhibiting HSV-1, 293T cells were transduced with HELP for 24 hr and infected with HSV-1. The supernatants were harvested 1 day and 2 days after HSV-1 infection, respectively, and subjected to virus yield assay. We found inhibitory effects for all viral gene-targeting mLPs with the UL8/UL29 co-targeting HELP the most efficient (Fig. 1d and Supplementary fig. 1). We therefore chose HELP in all the subsequent experiments.

**Figure 1.**
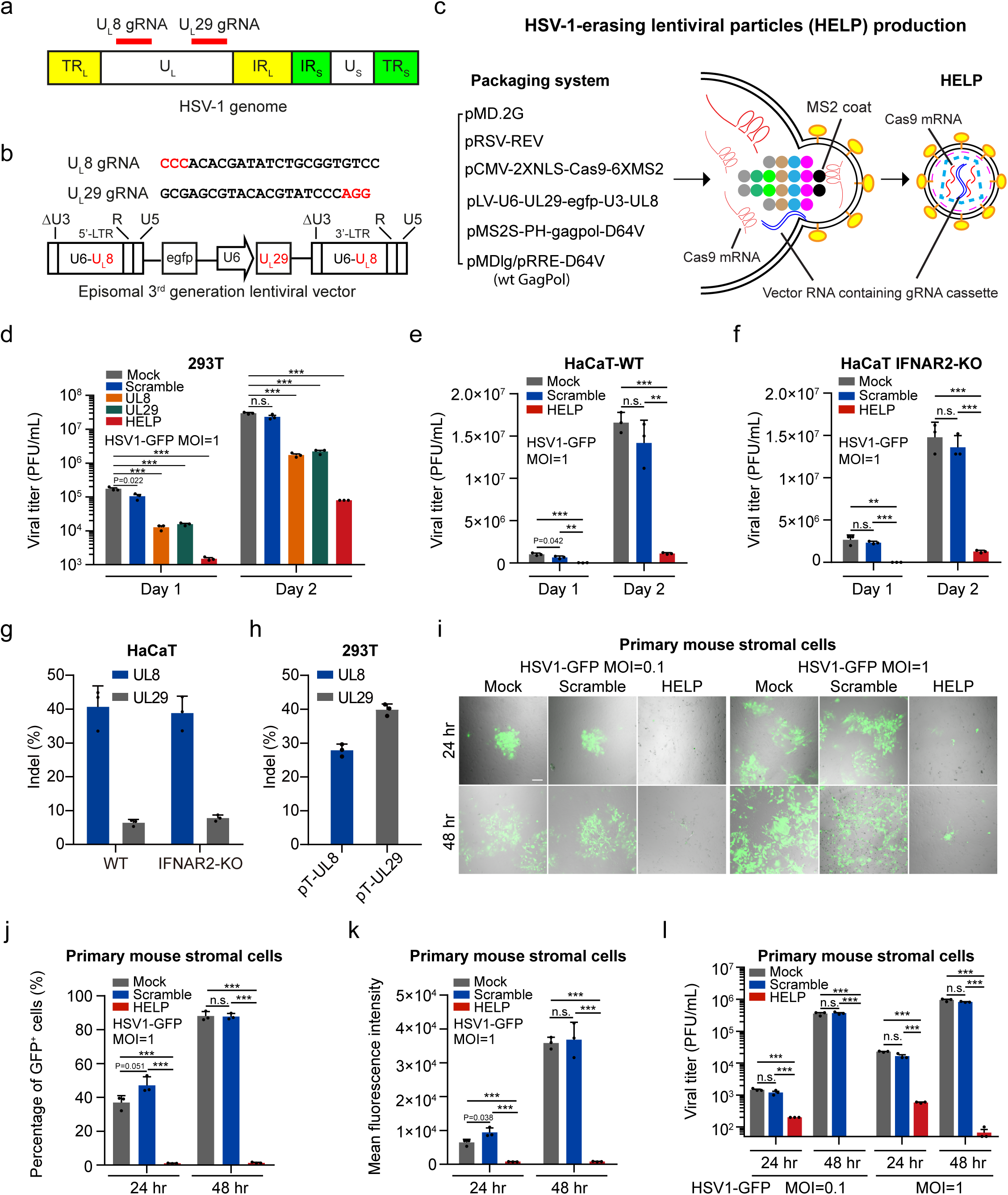
HELP blocks HSV-1 replication *in vitro*. **a**, Schematic representation of the HSV-1 genome. The long terminal repeats (TRL) and internal repeats (IRL), and the short internal (IRS) and terminal repeats (TRS) flanks the unique long (UL) and unique short (US) regions. The localization of targeted genes and gRNAs are indicated. **b**, The gRNA sequences and its expression cassette. Two gRNAs were cloned into the ΔU3 and the backbone of lentiviral vector, respectively, for HELP. For the single gRNA-carrying mLP, either UL8, UL29 or scramble gRNA was cloned into ΔU3. **c**, Schematic illustration of the packaging and production process. **d-f**, The anti-viral activity of HELP in 293T cells (**d**), wild-type HaCaT cells (**e**) and HaCaT IFNAR2-KO cells (**f**). Progeny viruses in the supernatants were titrated 1 day and 2 days after HSV-1 infection, respectively. **g**, TIDE analysis of indels induced by HELP in HSV-1 genome. Viral DNA was isolated from two-day samples in (**e**) and (**f**). **h**, TIDE analysis of indels induced by HELP in plasmids containing UL8 and UL29 target sequence (pT-UL8 and pT-UL29), respectively. **i-l**, The antiviral activity of HELP in primary mouse stromal cells evaluated by confocal microscopy (**i**), flow cytometry (**j** and **k**) and PFU analysis (**l**). Scale bars, 100 µm. Here, a GFP expressing strain HSV1-GFP was used. For all experiments, cells were seeded 24 hr before transduction (400 ng p24) at a density of 4×10^4^/well. 24 hr after transduction, cells were infected with HSV-1. Samples were harvested 1 day and 2 days after HSV-1 infection. n=3, error bars represent ±s.e.m. Unpaired two-tailed Student’s t-tests, **P< 0.01, ***P< 0.001, n.s.=non-significant.

The HSV-1 infection is sensitive to type I interferons (IFNs)induced by pathogen-associated molecular pattern molecules (PAMPs) even in the absence of gene editing (Supplementary fig. 2)^24^. To exclude the necessity of type I IFNs, here, we evaluated the antiviral activity of HELP in both wild-type and IFNAR2 knockout HaCaT cells. We found HELP significantly inhibited HSV-1 in both cell lines, but not the scramble control (Fig. 1e and Fig. 1f). Furthermore, we analysed the UL8 and UL29 loci and found indel frequency about 40% for UL8 while only 6% UL29 (Fig. 1g). The indels on UL29 were relatively low. ICP8 (encoded by UL29) plays a multifunctional role in the viral life cycle including DNA synthesis^25^. Therefore, mutations on UL29 are intolerant and quickly diluted. Indeed, when using plasmids as the target, we obtained even higher indels for UL29 gRNA (Fig. 1h). Notably, antiviral activity of HELP is under-estimated using PCR-based indels analysis, since not all the cleavage outcomes can be amplified (Supplementary fig. 3). In addition, we found HELP did not provoke innate immune sensing in contrast to HSV-1 strains which were all sensed by THP-1 derived macrophages at MOI=1 and induced moderate, but significant IFN response (Supplementary fig. 4). Together, these data suggest the HELP inhibits HSV-1 through DNA disruption, but not type I IFN-dependent innate immune response.

**Figure 2.**
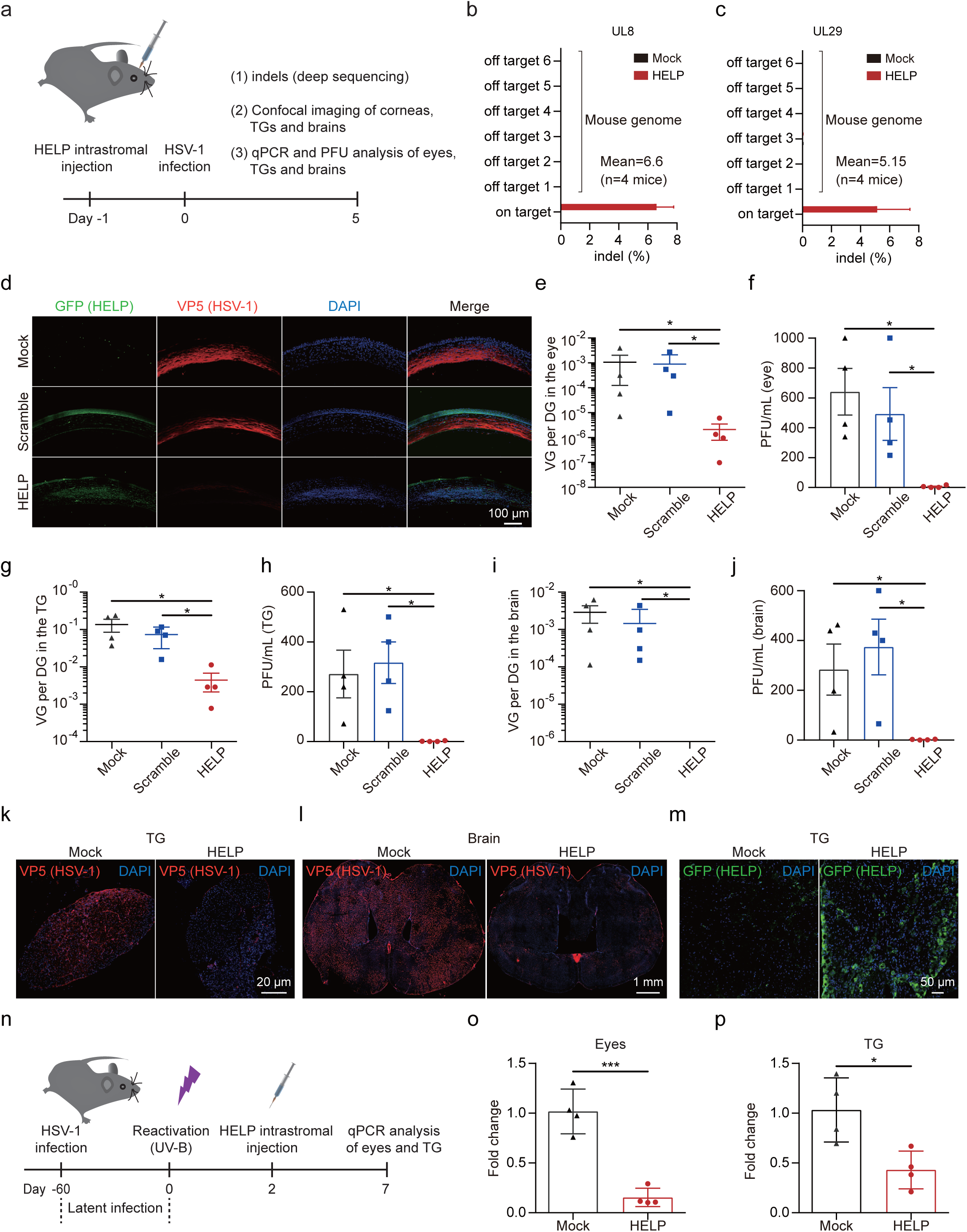
HELP blocks HSV-1 corneal and neuron infection *in vivo*. **a**, Flow chart for evaluating antiviral effects of HELP *in vivo*. 200 ng p24 HELP and scramble mLP, or 2 µL PBS were injected to corneas of C57/6J mice by intrastromal injection. After 24 hours, the mice were infected on both eyes with HSV-1 17syn+ (2×10^6^ PFU/eye). On day 5, all mice were sacrificed for corresponding analysis. **b** and **c**, Deep sequencing analysis of on-target effects in HSV-1 and off-target effects in mouse genome (n=4 mice). **d**, Confocal imaging of HSV-1 and HELP in corneas. Mouse corneal sections were incubated with both anti-GFP (HELP) and anti-HSV-1 (VP5) antibodies, respectively. Scale bars, 100 µm. **e-j**, qPCR and plaque-forming unit (PFU) analysis of HSV-1 dissemination in the eye, trigeminal ganglia (TG) and brains. The abundance of HSV-1 shown as viral genome (VG) per diploid genome (DG) (n=4 mice). **k** and **l**, Confocal analysis of HSV-1 presence in the TG and whole brain, respectively, after HELP treatment. Scale bars, 20 µm and 1 mm, respectively. **m**, Confocal analysis of HELP presence in the TG after intracorneal injection. Scale bars, 50 µm. **n**, Flow chart for evaluating antiviral activity using a recurrent HSK model. The mice were infected on both eyes with HSV-1 17syn+ (2×10^5^ PFU/eye) **o** and **p**, Virus load in the eyes and TG after HELP treatment of corneas. HSV-1 genome was detected by qPCR. Error bars represent ±s.e.m. Unpaired two-tailed Student’s t-tests, *P< 0.05, ***P< 0.001, n.s.=non-significant.

**Figure 3.**
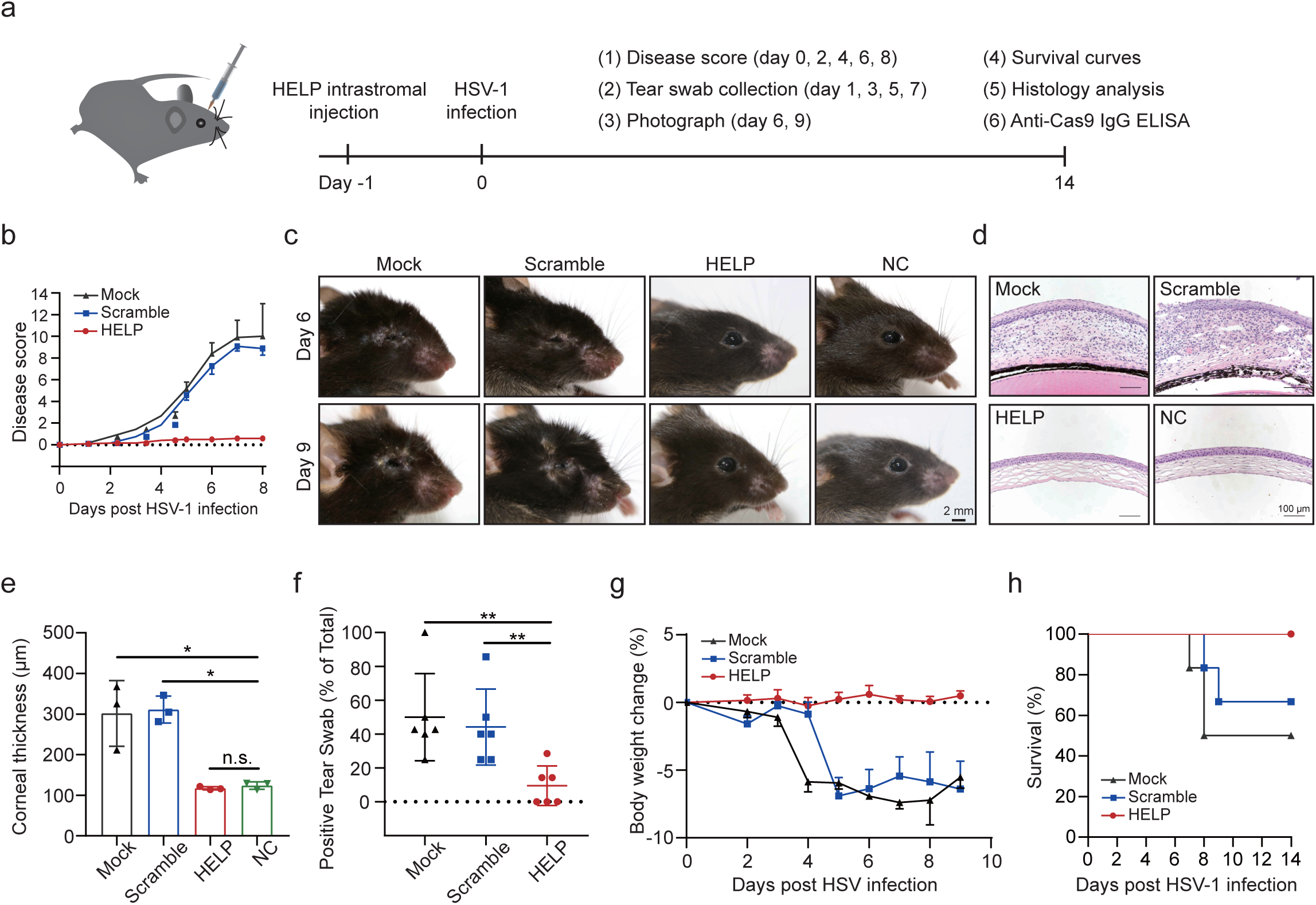
HELP suppresses HSV-1 associated disease pathologies. **a**, Flow chart for evaluating antiviral effects of HELP *in vivo*. 200 ng p24 HELP and scramble mLP, or 2 µL PBS were injected to corneas of C57/6J mice by intrastromal injection. After 24 hours, the mice were infected on both eyes with HSV-1 17syn+ (2×10^6^ PFU/eye). **b**, Ocular disease scores (0 to 4, 4 being severe) in mice (n=6 mice). **c**, Representative micrographs of the right eyes of differently treated mice on 6 dpi and 9 dpi. Scale bars, 2 mm. **d**, Representative corneal histology sections of eyes on 14 dpi. Scale bars, 100 µm. **e**, Thickness of the corneas assessed from histology (n = 3 mice). **f**, Secreted HSV-1 assessed from the swabs of eyes. Each mouse was collected for swabs at 1, 3, 5, 7 dpi during the experiment. Percentage of HSV-1 positive swabs was recorded (n=6 mice). **g**, Body weight (n=6 mice). **h**, Kaplan-Meier survival curves (n=6 mice). Error bars represent ±s.e.m. Unpaired two-tailed Student’s t-tests. *P< 0.05, ***P< 0.001, n.s.=non-significant.

**Figure 4.**
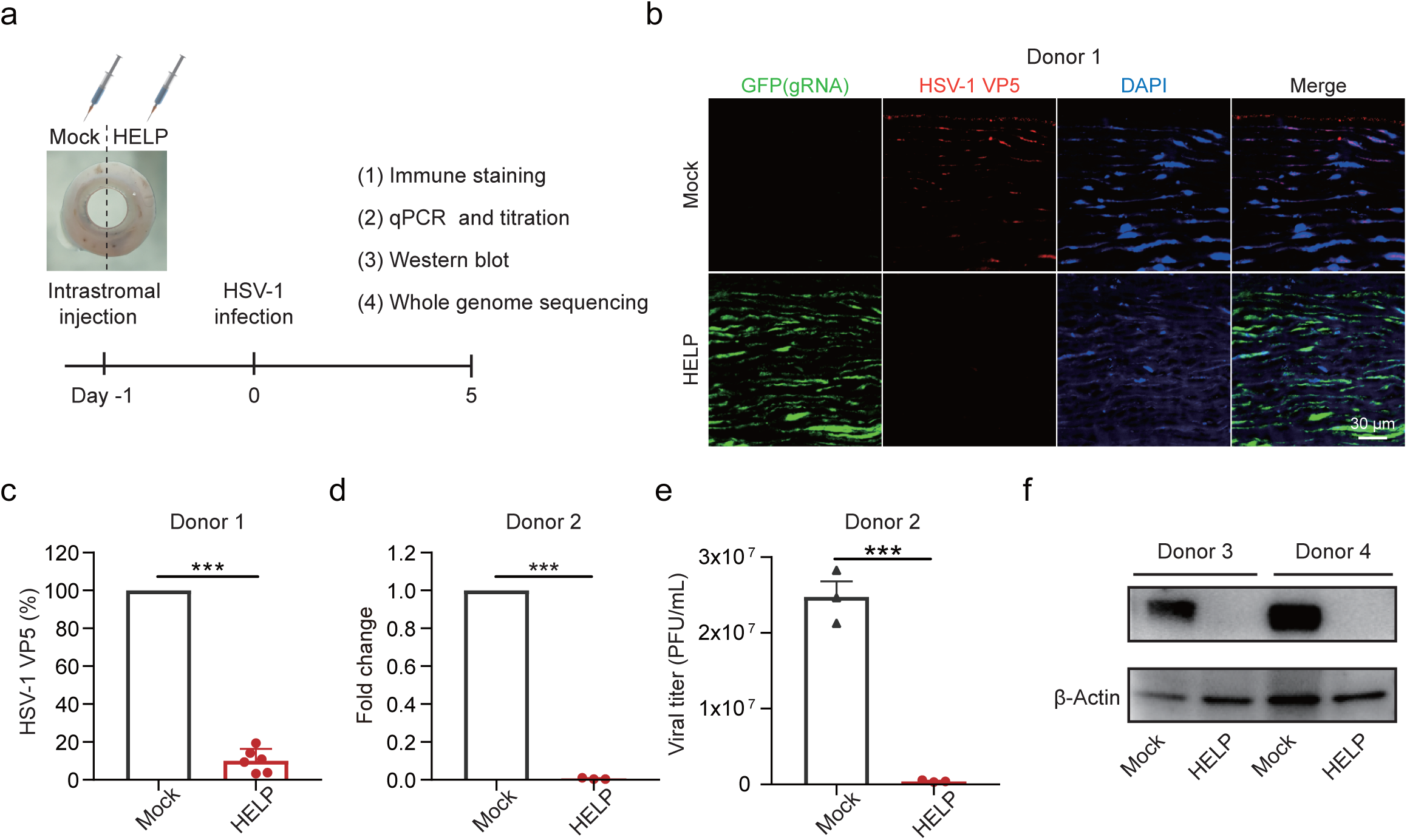
HELP eliminates HSV-1 in tissue culture of human corneas. **a**, Flow chart for evaluating antiviral effects of HELP in human corneas. 1.5 µg p24 HELP or 15 µL PBS were injected to the corresponding punch derived from the same human cornea. After 24 hours, the corneal punches were infected with HSV-1 17syn+ (2×10^6^ PFU/piece). **b**, Confocal analysis of the distribution of HSV-1 and HELP in the human cornea. GFP is indicative of the presence of HELP and VP5 (red) is indicative of HSV-1. Scale bar, 30 µm. **c**, Percentage of VP5+ cells presented in (**b**) (n = 6 sections) via Image J software and is represented as percentage of mock-treated tissues. **d**, qPCR analysis the fold change of HSV-1 genome in the human cornea after HELP treatment (n=3 pieces). **e**, Viral titers analysis of supernatants from human corneal cultures after HELP treatment (n=3 pieces). **f**, Western blot analysis of VP5 protein after HELP treatment. Error bars represent ±s.e.m. Error bars represent ±s.e.m. Unpaired two-tailed Student’s t-tests, ***P< 0.001.

Corneal stroma is highly linked to keratitis recurrence^26^. Stroma are rich of nerve terminals that originate from the trigeminal ganglia (TG) where HSV-1 maintains latancy^27^. Therefore, we explored if HELP was functional in primary corneal stromal cells from mice. The primary stromal cells were transduced with a non-GFP version HELP for 24 hr and then infected with HSV-1. We found HELP potently suppressed the GFP expression as well as viral replication using both low and high MOI on either 24 hr or 48 hr post-infection while the scramble control did not show any protection effects (Fig. 1i-l).

HSV-1 propagates fast (about 18 hr for the lytic replication cycle), so it could be expected that permanent nuclease presence is necessary to achieve therapeutic efficacy. Paradoxically, persisting nuclease expression may bring additional risks. From a safety perspective, transient nuclease exposure is desired for CRISPR therapeutics. Upon infection, HSV-1 encounters harsh antiviral response from innate and adaptive immunity. Here, we hypothesize that reducing the viral load to a certain level is sufficient to control the virus *in vivo*. To verify this, we performed dose-response experiments of HSV-1 infection on scarified corneas of mice. Indeed, only when HSV-1 is over 2×10^4^ PFU, did the viability and weight of mice change and symptoms of keratitis developed (Supplementary fig. 5).

We therefore set out to investigate the potential of HELP as a novel HSK therapeutic *in vivo*. To identify the kinetics of HSV-1 infection in our HSK model, we visualized HSV-1 using confocal imagining and found the virus progressively disseminated from surficial to deep side of corneal stroma during the time from day 2 to 8 post-infection (Supplementary fig. 6). We therefore chose day 5 as our endpoint. The experimental set-up is illustrated in Fig. 2a. We first deep-sequenced the on-target activity of HELP on HSV-1 genome in murine corneas. The indels induced by HELP was approximately 7% on UL8 loci and 5% for UL29 loci (Fig. 2b). Next, we performed confocal imaging to access HSV-1 replication and HELP distribution in the corneas of mice, which was represented by viral capsid protein VP5 and GFP, respectively. We found HSV-1 was replicating actively in the corneal stroma in the mock and scramble controls while it was hardly detectible after HELP treatment (Fig. 2c). Accordingly, HELP was evenly distributed in all corneal structures-from epithelium and stroma to endothelium (Fig. 2c). To assess whether HELP treatment blocks the transmission of HSV-1 from corneal epithelium to the peripheral and central nervous system (CNS), eyes, TGs and brain samples from all the infected mice were harvested and examined for the copy number of HSV-1 genomes and infectious viruses. In all samples, the viral load was significantly reduced after HELP treatment (Fig. 2e-2j). Additionally, we performed confocal imaging of TG and whole brain. In agreement with the qPCR and PFU analysis, we found HELP diminished HSV-1 to almost undetectable level in both TG and brain (Fig. 2k-2l). Tissue distribution is an important safety index for *in vivo* gene therapy. We therefore evaluated the dissemination of HELP in the whole body and found HELP was highly restricted to the eyes, not other organs, including reproductive organs (Supplementary fig. 7). Interestingly, we also detected HELP in the TG though it was factually injected in the corneas supporting retrograde delivery of CRISPR from neuron termini in the corneas to neuron body in the TG (Supplementary fig. 7). This finding was further strengthened detection of HELP in the TG by confocal imaging (Fig. 2m).

To mimic the natural disease process of HSK, we adopted a recurrent HSK model in which eyes were infected with HSV-1 to establish latency before of HELP treatment (Fig. 2n). We reactivated the mice survived from the acute infection by UV-B irradiation of the eyes 60 days after HSV-1 inoculation. We then treated the eyes with HELP and quantified the HSV-1 genome two days later. Indeed, we found HELP significantly decreased viral DNA in the infected eyes (Fig. 2o). Notably, we also found reduced level of HSV-1 reservoir in the TG indicating HELP was able to modulate the viral reservoir in the TG (Fig. 2p).

To determine the disease development and therapeutic efficacy, we monitored the clinical signs of acute ocular herpes infection and scored them in a blinded fashion (Fig. 3a). Mice that were treated with HELP did not show any disease progress (n=6 mice) while the mock-treated and scramble gRNA treated eyes developed severe signs of ocular infection (Fig. 3b and Fig. 3c). Next, we performed histological staining to examine the pathology of the eye. The mock- and scramble gRNA-treated eyes presented with irregular stroma matrix and increased corneal thickness, typical of acute infection (Fig. 3d and 3e). We further found HSV-1 infection in the corneas induced significant type I IFN response, while HELP transduction was not sensed (Supplementary Fig. 8). T-cell mediated destruction is responsible for much of the morbidity in infectious stromal disease in addition to direct viral effects^28^. Using immunohistochemistry, we showed HSV-1 infection leaded to CD4^+^ and CD8^+^ cells infiltration in the corneal stroma for the mock and the scramble groups, but HELP treatment prevented immune cell infiltration (Supplementary Fig. 9). We also noticed that PD-L1 was up-regulated in the epithelium and stroma of untreated mice after HSV-1 infection in consistent with previous observations (Supplementary Fig. 9)^29^. Increased local PD-L1 expression may inhibit virus clearance by immune cells highlighting the importance of direct DNA degradation by CRISPR. To access the secreted virus, the swabs were titrated every other day post-infection. HELP treatments significantly reduced the viral presence in the eyes (Fig. 3f). In addition, body weights were recorded every other day from 2 dpi. No loss of body weights was observed for HELP treated mice, while it was evident for the mock and scramble control (Fig. 3g). Survival curves were generated for each group to assess the efficacy of the treatments. Notably, all mice survived in the HELP treated groups (Fig. 3h). Finally, we examined if intrastromal injection of HELP induces Cas9-specific IgG in the blood stream. We did not observe significantly higher Cas9-specific IgG for both HELP and scramble control than the mock (n=5 mice, non-significant, Student’s t-tests). In contrast, when HELP was injected via footpad route, it provoked significantly higher anti-Cas9 IgG in the sera (n=5 mice, P<0.001, Student’s t-tests) (Supplementary Fig. 10). Taken together, these results suggest that the administration of HELP significantly reduced the manifestation of disease severity during ocular HSV-1 infection.

Having established the therapeutic efficacy of HELP in murine model, we sought to investigate the antiviral potential of HELP in human corneas (Fig. 4a). One human cornea was evenly divided into two halves, and injected with either 15 µL HELP (equal to 1.5 µg p24) or PBS, respectively, for confocal imaging. We found HELP was evenly spread in the stroma and potently inhibit HSV-1 replication as manifested by viral protein VP5 compared to mock controls (Fig. 4b-c). With cornea from another donor, we found HELP treatment significantly diminished both genome of HSV-1 and viral titers (Fig. 4d-e). Additionally, we showed that VP5 protein was hardly detectible after HELP treatment by Western blot analysis of corneal samples (Fig. 4f). Altogether, our data indicate that HELP efficiently inhibits HSV-1 replication in human corneas from four different donors.

Recently, Jaishankar et al. reported BX795-a commonly used inhibitor of TANK-binding kinase 1 (TBK1)-blocks HSV-1 infection *in vivo* by targeting Akt phosphorylation in infected cells^30^. However, BX795 does not eliminate viral DNA and 20% of treated-mice still died within 14 days. Using a similar model, all mice survived after HELP treatment though the death rate was even higher for controls.

HELP carries Cas9 in the form of mRNA and has no long-term off-targeting risk troubling other means of CRISPR delivery such as AAV. We show ‘hit-and-run’ gene editing is sufficient to achieve therapeutic efficacy against HSK *in vivo*, and block HSV-1 replication in human corneas. Additionally, HELP gRNAs targets to the genome of virus instead of human, which may accelerate it for clinical translation. Modulating virus reservoir is essential to prevent HSK from recurrence. Our study showed preliminary evidence of retrograde transportation of HELP from corneas to TG using VSV-G pseudotyping, however, enhanced retrograde delivery may be achieved by pseudotyping HELP with derivatives of rabies virus glycoprotein, which may further strengthen the efficacy of HELP in clinical studies^31,32^.

## METHODS

### Cell cultures and HSV-1 propagation

293T, Vero, HaCaT and HaCaT IFNAR2-KO cells were cultured in DMEM (Gibco, USA). DMEM media were supplemented with 10% fetal bovine serum (Gibco, USA) and 1% penicillin/streptomycin (P/S) (Thermo Fisher Scientific, USA). Primary mouse corneal stromal cells were maintained in MEM (Gibco, USA) supplemented with 1% P/S and 10% fetal bovine serum. Human cornea tissues were maintained in MEM (Gibco, USA) supplemented with 1% P/S and 10% fetal bovine serum. All cells were cultured at 37°C and 5% (vol/vol) CO_2_. HSV1 strain 17syn+, mckrea, F, and HSV1-GFP were propagated in and titrated on Vero cells.

### Production of mLPs

293T cells were seeded in 15-cm dishes at a density of 10^7^/dish 24 hr before calcium phosphate transfection. 24 hr after transfection, the media were refreshed, and the supernatants were harvested 48 hr and 72 hr post-transfection before passing through a 0.22-µm filter (Millipore, USA) and ultracentrifugation at RPM 25,000 at 4°C for 2 hr. Pellets were re-suspended in PBS and stored at -80°C. To produce ‘all-in-one’ mLPs, 293T cells were transfected with 9.07 µg pMD.2G, 7.26 µg pRSV-Rev, 15.74 µg pMDlg/pRRE-D64V, 15.74 µg pMS2M-PH-gagpol-D64V, 31.46 µg pCMV-Cas9-6XMS2 and 31.46 µg pLV-egfp-U3-osp-gRNA with corresponding gRNA sequence. To produce HELP, pLV-U6-UL29-egfp-U3-UL8 was used as gRNA producing plasmid. To produce non-GFP version HELP, pLV-U6-UL29-U3-UL8 was used. The gene sequences of plasmids were supplied.

### HSV-1 plaque assay

HSV-1 plaque assays were performed in triplicates for each biological sample. 1.5×10^5^ Vero cells were seeded in a 12-well plate in the complete DMEM and infected the following day with various dilutions of HSV stocks or culture supernatants. Two hours after infection, cells were overlaid with 1% agarose (Sangon, China) solution. After incubation for 3 days, cells were fixed with 4% formaldehyde and stained using 1% crystal violet solution at room temperature for 2 hours. After 3 washes with PBS, plates were allowed to dry and the numbers of plaques were counted. Viral titers were calculated as plaque forming units/mL (PFU/mL).

### Infection of cells

For 293T, HaCaT, HaCaT-IFNAR2-KO, THP1 and primary mouse corneal stromal cells, 4×10^4^ cells were seeded in a 48-well plate well and transduced with 400 ng mLP on the following day. The media were refreshed 12 hpi. After 24-hr transduction, the cell infected with HSV1-GFP at a MOI of 1. The supernatants were harvested at 24 and 48 hpi for plaque assay, and the DNA isolated from the cell lysates using viral DNA extraction kit (TaKaRa, Japan) was used to determine the cleavage activity of HSV-1 genomes by Sanger sequencing and TIDE analysis. The used primers are shown in the Supplementary Table 1.

### Flow cytometry analysis

The primary mouse stromal cells were seeded at a density 4×10^4^/well on day 1, and transduced with UL29-UL8-co-targeting mLP-CRISPR (non-GFP version) on day 2. The cells were then infected with HSV1-GFP on day 3. The GFP signals were determined by flow cytometry (BD & LSR Fortessa) and analysed for the percentage of GFP^+^ cells and mean fluorescence intensity on day 4 and 5.

## ELISA

The p24 of mLP was measured using HIV p24 ELISA according to the manufacturer (Beijing Biodragon Immunotechnologies, China). To detect mouse humoral IgG immune response to Cas9, IgG Mouse ELISA Kit (Abcam, USA) was uses following manufacturer’s protocol with a few modifications. A total of 0.25 µg of recombinant Cas9 proteins (Novoprotein, China) suspended in PBS were used to coat 96-well ELISA plates and incubated at 4°C for 12 hr, then washed three times using 1X Wash Buffer. Plates were blocked with 2% BSA Blocking Solution for 2 hr at room temperature, then washed three times. Serum samples were added to each well. The remaining steps were doing as manufacturer’s protocol. Cas9 Mouse mAb (Cell Signaling Technology, USA) was used to make a standard curve, the dilution gradient was accorded to the instruction of IgG Mouse ELISA Kit.

### Western blotting

To detect the effect of HSV infected human stromal cornea after treatment with UL29-UL8-targeting mLP-CRISPR. The human cornea stromal tissues were grinded using Tissuelyser with magnetic beads. The suspensions were lysed in RIPA (Beyotime Biotechnology, China) in the presence of a protease inhibitor (Beyotime Biotechnology, China) for 30 minutes and incubated with SDS-PAGE sample loading buffer (Beyotime Biotechnology, China) for 15 minutes at 98°C. The proteins were separated by SDS-polyacrylamide gel electrophoresis and transferred to the PVDF membrane. Membrane was blocked by 5% fat-free milk dissolved in TBS/0.05% Tween-20 for 1 hr. Membrane was cut off according marker and incubated with primary antibody overnight at 4°C. The membranes were incubated with anti-mouse secondary antibodies (Cell Signaling Technology, USA) and visualized by hypersensitive ECL chemiluminescence (Beyotime Biotechnology, China). Actin was used for signal normalization across samples. The primary antibodies used in this experiment were HSV VP5 monoclonal antibody (Santa Cruz Biotechnology, USA) and β-Actin Mouse mAb (Cell Signaling Technology, USA).

### Quantitative PCR

Genomic DNAs and total RNAs from all samples were extracted using Viral DNA/RNA extraction kit (TaKaRa, Japan). cDNA was synthesized using the QuantScript RT Kit (TIANGEN, China) according to the manufacturer’s protocol. RT-qPCR was performed using PowerUp SYBR Green Master Mix (Applied Biosystems, USA) following the manufacturer’s protocol. For HSV-1 genome quantification in mouse eye, trigeminal ganglion and brain, genomic DNAs were extracted from the eye, trigeminal ganglion and brain. qPCR was performed to detect HSV-1 (primer Y5/Y6) which was then normalized to GAPDH (SK13/SK14). To detect HSV-1 relative expression in human cornea and mouse model of recurrent HSK, total RNA was extracted. RT-qPCR was performed using primers Y5/Y6, then normalized to human GAPDH (SK55/SK56) or mouse GAPDH (SK13/SK14). To detect mLP distribution in vivo, genomic DNAs were extracted from eye, trigeminal ganglion, heart, liver, spleen, lung, kidney and testis. qPCR was performed to detect WPRE (primer SK9/SK10) which was then normalized to GAPDH (SK13/SK14). To detect immunoreaction induced by HSV1 in mouse, total RNA was extracted from eyes. RT-qPCR was performed to detected ISG15 (SK51/SK52), RIG1 (Y7/Y8) and IFNB1 (Y9/Y10) which was then normalized to GAPDH (SK13/SK14). To detect immunoreaction induced by HSV1 in cells THP1 and induced by HELP in cells HaCat, total RNA was extracted. RT-qPCR was performed to detected ISG15 (Y11/Y12), RIG1 (Y13/Y14) and IFNB1 (Y15/Y16) which was then normalized to GAPDH (SK55/SK56). The used primer sequences were listed in the Supplementary Table 2.

### Mice

6-8 weeks old, male, specific-pathogen-free (SPF) C57BL/6J mice were used in this study. The mLP or PBS was injected to mice by intrastromal injection under the dissecting microscope. All mouse in this study were complied with the guidelines of the Institutional Animal Care and Use Committee (IACUC) of the Shanghai Jiao Tong University.

### Intrastromal injection

The mice were anesthetized by intraperitoneal injection of 1.25% Avertin (0.2 mL/10 g of body weight). A small intrastromal pocket was carefully created in the mid-peripheral cornea by 29 G needle. Then a 33G syringe was inserted towards the central cornea, and 2 µL mLP or PBS was injected to corneal stroma. Both eyes were injected in this study.

### Murine acute HSV-1 infection model

The mice were anesthetized by intraperitoneal injection of 1.25% Avertin. Corneas were scarified with a 3×3 crosshatch pattern. The mice were inoculated with 2×10^6^ PFU HSV-1 17syn+ on both eyes. The body weight and disease scores were measured at the indicated times post infection. The scoring was performed as blinded study: hair loss (0: none, 1: minimal periocular hair loss, 2: moderate periocular hair loss, 3: severe hair loss limited to periocular, 4: hair loss severe and extensive); hydrocephalus (0: none, 1: minor bump, 2: moderate bump, 3: large bump); symptoms related to neurological disease (0: normal, 1: jumpy, 2: uncoordinated, 3: hunched/lethargic, 4: unresponsive/no movement); eye swell/lesions (0: none, 1: minor swelling, 2: moderate swelling, 3: severe swelling and skin lesions, 4: lesions extensive). Mice were sacrificed at the specified times post infection. To collect the eye swabs, mouse eyes were gently proptosed, then wiping a sterile cotton swab (Miraclean Technology, China) three times around the eye in a circular motion and twice across the center of the cornea in an “+” pattern. The cotton swabs were placed in 1 mL of DMEM containing 2% (vol/vol) FBS, 1% P/S and stored at -80°C until titrated by plaque assay. Serum were collected at 14 dpi to test the mouse humoral IgG immune response to Cas9 by ELISA. Three mouse eyes from each group were processed for histological evaluation.

### Murine recurrent HSV-1 infection model

The mice were inoculated with 2×10^5^ PFU HSV-1 17syn+ for both eyes on scarified corneas. Mice survived from acute infection were maintained for 60 days and reactivated by UV-B irradiation of the eyes, followed by HELP treatment. The TGs and eyes were collected to quantify HSV-1 DNA by qPCR.

### Human cornea HSV-1 infection

The human corneas were obtained from fresh cadavers and supplied by Eye, Ear, Nose and Throat Hospital, Fudan University with experiments conducted according to the Declaration of Helsinki and in compliance with China law. The corneas were evenly divided into two halves. One half was dosed with 15 µL HELP by intrastromal injection (equal to 1.5 µg p24) while the other was dosed with PBS by intrastromal injection as a mock control. The corneas were then infected with 2×10^6^ PFU of HSV-1 17syn+ in MEM medium containing 2% FBS, and the media were refreshed 2 hpi with MEM containing 10% FBS and 5% P/S. Two days after HSV-1 infection, the corneas were processed for immunofluorescence imaging, immunoblotting or DNA isolation by the viral DNA extraction kit to determine the viral genomes by qPCR. The supernatants were collected for plaque assay.

### Immunofluorescence imaging

For confocal imaging, the 293T and mouse cornea stroma cell were imaged under A1Si Laser Scanning Confocal Microscope (Nikon, Japan) at the indicated times. The eyes, trigeminal ganglions and brains were fixed in 4% PFA overnight at 4°C then transfer to 30% sucrose until the tissue sink. All tissues were embedded in OCT Tissue-Tek (Sakura Finetek, USA) and frozen on liquid nitrogen. Sections with 10-µm thickness were cut using the CM1950 (Leica, Germany) freezing microtome and processed for immunofluorescence as mentioned previously. The slides were dried at room temperature for 10 minutes and blocked in blocking buffer with 5% normal donkey serum (Solarbio, China), 1% BSA, 0.3% Triton X-100 in PBS in humidified box at room temperature for 30 minutes. The slides were incubated with the primary antibody against HSV-1 VP5 (Santa Cruz Biotechnology, USA) or GFP (GeneTex, USA) in 1% BSA overnight at 4°C. After washing, the slides were incubated with a secondary antibody (Santa Cruz Biotechnology, USA) in 1% BSA for 1 hour.

### Deep sequencing

The top 5 or 6 predicted off-target sites in human and mouse genome for UL29-targeting and UL8-targeting gRNA were identified by Cas-OFFinder online predictor respectively. The on-target and predicted off-target regions were PCR amplified and were pooled by an equal molar ratio for double-end sequencing using Illumina MiSeq at Novogene. Raw data of Next-generation sequencing were analysed by online Cas-analyzer. The primer sequences are listed in the Supplementary Table 2.

### Histology

Mouse eyes were dissected and fixed in paraformaldehyde before embedding in paraffin, sectioning at 10-µm thickness and staining with hematoxylin and eosin. For immunohistochemistry staining, the sections were de-paraffinated and rehydrated followed by incubated with citrate buffer for antigen retrieval. To block endogenous peroxidase activity, the sections were treated by 3% H_2_O_2_ for 25 minutes. The sections were then blocked with 3% BSA at room temperature for 30 minutes, followed by incubating with anti-CD4+, anti-CD8+ and anti-PD-L1 antibody at 4°C overnight. The slides were then incubated with a secondary antibody, followed by incubating with the freshly prepared DAB substrate solution to reveal the color of antibody. At last, the tissue slides were counterstained with hematoxylin and blued with ammonia water, and then dehydrated and coverslipped.

## Statistics

Data are presented as mean ± s.e.m. in all experiments (n≥3). Student’s t-tests were performed to determine the P values. The specific statistical method applied, description of replicates can be found in the figure legends. *indicates statistical significance (*P< 0.05, **P< 0.01, ***P< 0.001, n.s.=non-significant).

## Data availability

Data generated or analysed during this study are available from the corresponding author on reasonable request.

## Acknowledgement

J.H. is supported by the National Natural Science Foundation of China [81970766 and 81670818] and the Shanghai Rising-Star Program [18QA1401100]. Y.C. is supported by National Natural Science Foundation of China [31971364], Pujiang Talent Project of Shanghai [GJ4150006], Shanghai Municipal Natural Science Foundation [BS4150002] and Startup funding from Shanghai Center for Systems Biomedicine, Shanghai Jiao Tong University [WF220441504].

## Author contribution

D.Y, S.L., J.H. and Y.C. conceived the study and designed the experiments; D.Y, S.L., D.W., Y.D. and H.J performed the experiments; all the authors analysed the data; Y.D., S.L. and Y.C. wrote the manuscript with the help from all the authors.

## REFERENCES

1. Liesegang, T.J. Herpes simplex virus epidemiology and ocular importance. Cornea 20, 1–13 (2001).

2. Paludan, S.R., Bowie, A.G., Horan, K.A. & Fitzgerald, K.A. Recognition of herpesviruses by the innate immune system. Nat. Rev. Immunol. 11, 143–154 (2011).

3. Bradshaw, M.J. & Venkatesan, A. Herpes Simplex Virus-1 Encephalitis in Adults: Pathophysiology, Diagnosis, and Management. Neurotherapeutics 13, 493–508 (2016).

4. Farooq, A.V. & Shukla, D. Herpes simplex epithelial and stromal keratitis: an epidemiologic update. Surv. Ophthalmol. 57, 448–462 (2012).

5. Vadlapudi, A.D., Vadlapatla, R.K. & Mitra, A.K. Update on emerging antivirals for the management of herpes simplex virus infections: a patenting perspective. Recent Pat Antiinfect Drug Discov 8, 55–67 (2013).

6. Chilukuri, S. & Rosen, T. Management of acyclovir-resistant herpes simplex virus. Dermatol. Clin. 21, 311–320 (2003).

7. Jiang, Y.C., Feng, H., Lin, Y.C. & Guo, X.R. New strategies against drug resistance to herpes simplex virus. Int. J. Oral Sci. 8, 1–6 (2016).

8. Lass, J.H., Langston, R.H., Foster, C.S. & Pavan-Langston, D. Antiviral medications and corneal wound healing. Antiviral Res. 4, 143–157 (1984).

9. Moshirfar, M., et al. A Review of Corneal Endotheliitis and Endotheliopathy: Differential Diagnosis, Evaluation, and Treatment. Ophthalmol Ther 8, 195–213 (2019).

10. Hillenaar, T., Weenen, C., Wubbels, R.J. & Remeijer, L. Endothelial involvement in herpes simplex virus keratitis: an in vivo confocal microscopy study. Ophthalmology 116, 2077-2086 e2071-2072 (2009).

11. Tucker, W.E., Jr., et al. Preclinical toxicology studies with acyclovir: ophthalmic and cutaneous tests. Fundam. Appl. Toxicol. 3, 569–572 (1983).

12. Jayamanne, D.G., Vize, C., Ellerton, C.R., Morgan, S.J. & Gillie, R.F. Severe reversible ocular anterior segment ischaemia following topical trifluorothymidine (F3T) treatment for herpes simplex keratouveitis. Eye (Lond) 11 (Pt 5), 757–759 (1997).

13. Spiegal, D.M. & Lau, K. Acute renal failure and coma secondary to acyclovir therapy. JAMA 255, 1882–1883 (1986).

14. Yildiz, C., Ozsurekci, Y., Gucer, S., Cengiz, A.B. & Topaloglu, R. Acute kidney injury due to acyclovir. CEN Case Rep 2, 38–40 (2013).

15. Koganti, R., Yadavalli, T. & Shukla, D. Current and Emerging Therapies for Ocular Herpes Simplex Virus Type-1 Infections. Microorganisms 7, 429 (2019).

16. Nelson, C.E., et al. Long-term evaluation of AAV-CRISPR genome editing for Duchenne muscular dystrophy. Nat. Med. 25, 427–432 (2019).

17. Maeder, M.L., et al. Development of a gene-editing approach to restore vision loss in Leber congenital amaurosis type 10. Nat. Med. 25, 229–233 (2019).

18. Beyret, E., et al. Single-dose CRISPR-Cas9 therapy extends lifespan of mice with Hutchinson-Gilford progeria syndrome. Nat. Med. 25, 419–422 (2019).

19. Santiago-Fernandez, O., et al. Development of a CRISPR/Cas9-based therapy for Hutchinson-Gilford progeria syndrome. Nat. Med. 25, 423–426 (2019).

20. Lee, B., et al. Nanoparticle delivery of CRISPR into the brain rescues a mouse model of fragile X syndrome from exaggerated repetitive behaviours. Nat Biomed Eng 2, 497–507 (2018).

21. Gao, X., et al. Treatment of autosomal dominant hearing loss by in vivo delivery of genome editing agents. Nature 553, 217–221 (2018).

22. de Buhr, H. & Lebbink, R.J. Harnessing CRISPR to combat human viral infections. Curr. Opin. Immunol. 54, 123–129 (2018).

23. van Diemen, F.R., et al. CRISPR/Cas9-Mediated Genome Editing of Herpesviruses Limits Productive and Latent Infections. PLoS Pathog. 12, e1005701 (2016).

24. Reinert, L.S., et al. Sensing of HSV-1 by the cGAS-STING pathway in microglia orchestrates antiviral defence in the CNS. Nat. Commun. 7, 13348 (2016).

25. Weller, S.K. & Coen, D.M. Herpes simplex viruses: mechanisms of DNA replication. Cold Spring Harb. Perspect. Biol. 4, a013011 (2012).

26. Oral acyclovir for herpes simplex virus eye disease: effect on prevention of epithelial keratitis and stromal keratitis. Herpetic Eye Disease Study Group. Arch. Ophthalmol. 118, 1030–1036 (2000).

27. Kennedy, D.P., et al. Ocular herpes simplex virus type 1: is the cornea a reservoir for viral latency or a fast pit stop? Cornea 30, 251–259 (2011).

28. Newell, C.K., Martin, S., Sendele, D., Mercadal, C.M. & Rouse, B.T. Herpes simplex virus-induced stromal keratitis: role of T-lymphocyte subsets in immunopathology. J. Virol. 63, 769–775 (1989).

29. Jeon, S., Rowe, A.M., Carroll, K.L., Harvey, S.A.K. & Hendricks, R.L. PD-L1/B7-H1 Inhibits Viral Clearance by Macrophages in HSV-1-Infected Corneas. J. Immunol. 200, 3711–3719 (2018).

30. Jaishankar, D., et al. An off-target effect of BX795 blocks herpes simplex virus type 1 infection of the eye. Sci. Transl. Med. 10, eaan5861 (2018).

31. Kobayashi, K., et al. Pseudotyped Lentiviral Vectors for Retrograde Gene Delivery into Target Brain Regions. Front Neuroanat 11, 65 (2017).

32. Kato, S., et al. Enhancement of the transduction efficiency of a lentiviral vector for neuron-specific retrograde gene delivery through the point mutation of fusion glycoprotein type E. J. Neurosci. Methods 311, 147–155 (2019).

