## Supplementary Figures for "Intracorneal delivery of HSV-targeting CRISPR/Cas9 mRNA prevents herpetic stromal keratitis"

**Table of Contents**

**Supplementary Figures**

**Supplementary_Fig_S1.** The antiviral activity of HELP in 293T cells.

**Supplementary_Fig_S2.** Innate immune stimulator inhibits HSV1-GFP replication.

**Supplementary_Fig_S3.** Illustration of the possible outcomes of HSV-1 genome after HELP cleavage.

**Supplementary_Fig_S4.** Innate immune response induced by HELP and different strains of HSV-1 in THP-1 derived macrophages.

**Supplementary_Fig_S5.** Dose-response and keratitis symptoms.

**Supplementary_Fig_S6.** Time course of HSV-1 infection in the corneas.

**Supplementary Figure_S7.** Tissue distribution of HELP in the whole body.

**Supplementary_Fig_S8.** Innate immune response induced HELP and HSV-1 infection *in vivo*.

**Supplementary_Fig_S9.** T cell infiltration and PD-L1 expression in the corneas *in vivo*.

**Supplementary_Fig_S10.** Cas9-specific IgG in the serum.

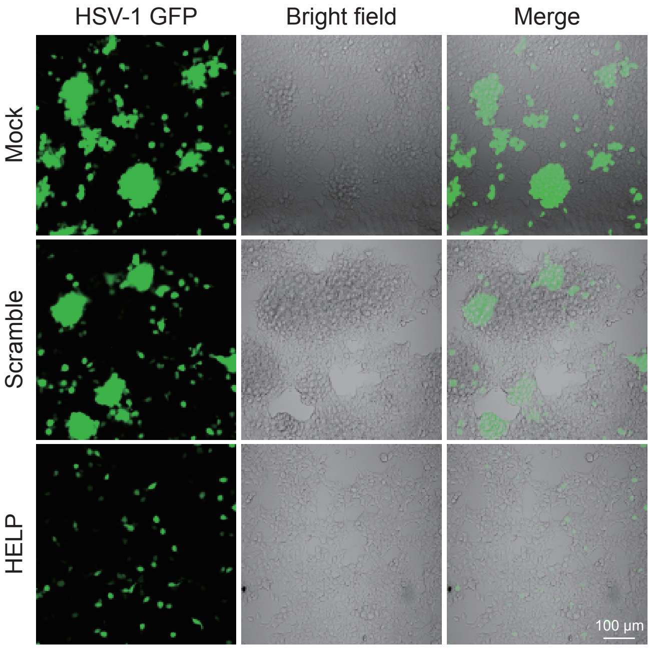


**Supplementary Figure 1. The antiviral activity of HELP in 293T cells.** Cells were seeded 24 hr before transduction of HELP (400 ng p24) at a density of $4\times10^4$/well. 24 hr after transduction, cells were infected with HSV1-GFP. Photographing was performed two days after HSV1-GFP infection (MOI=1).

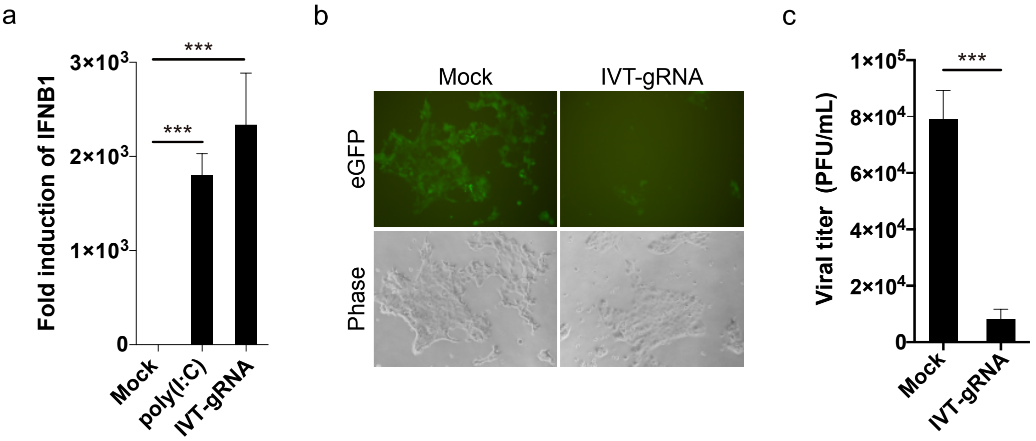


**Supplementary Figure 2. Innate immune stimulator inhibits HSV1-GFP replication. a**, Fold induction of IFNB1 by UL8-targeting *in vitro* transcribed gRNA (IVT-gRNA) and poly(I:C) in HaCaT cells. RNA was isolated from HaCaT cells 6 hr after infection or transfection. **b**, The antiviral activity of IVT-gRNA. Representative fluorescent and phase contrast photographs of HaCaT cells 24 hr after of IVT-gRNA transfection. IVT-gRNA transfection was performed 1 hr after HSV1-GFP infection (MOI=1.5). No Cas9 was provided. **c**, Plaque assay analysis of infectious viruses in supernatants harvested from mock and IVT-gRNA treated cells (from **b**). Error bars represent ±s.e.m. Unpaired two-tailed Student's t-tests were performed, ****P*< 0.001.

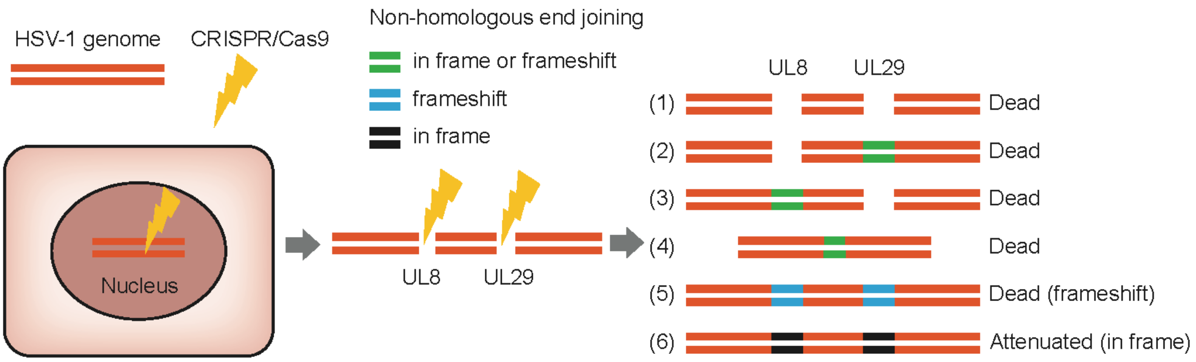


**Supplementary Figure 3. Illustration of the possible outcomes of HSV-1 genome after HELP cleavage.** If one of the two DSBs is not repaired (outcome 1-3), the viral genome un-replicable. If the breaks are repaired but causing large deletion or frameshift, the virus is also dead (outcome 4 and 5). If the DSBs are repaired and in frame, the virus is alive but attenuated.

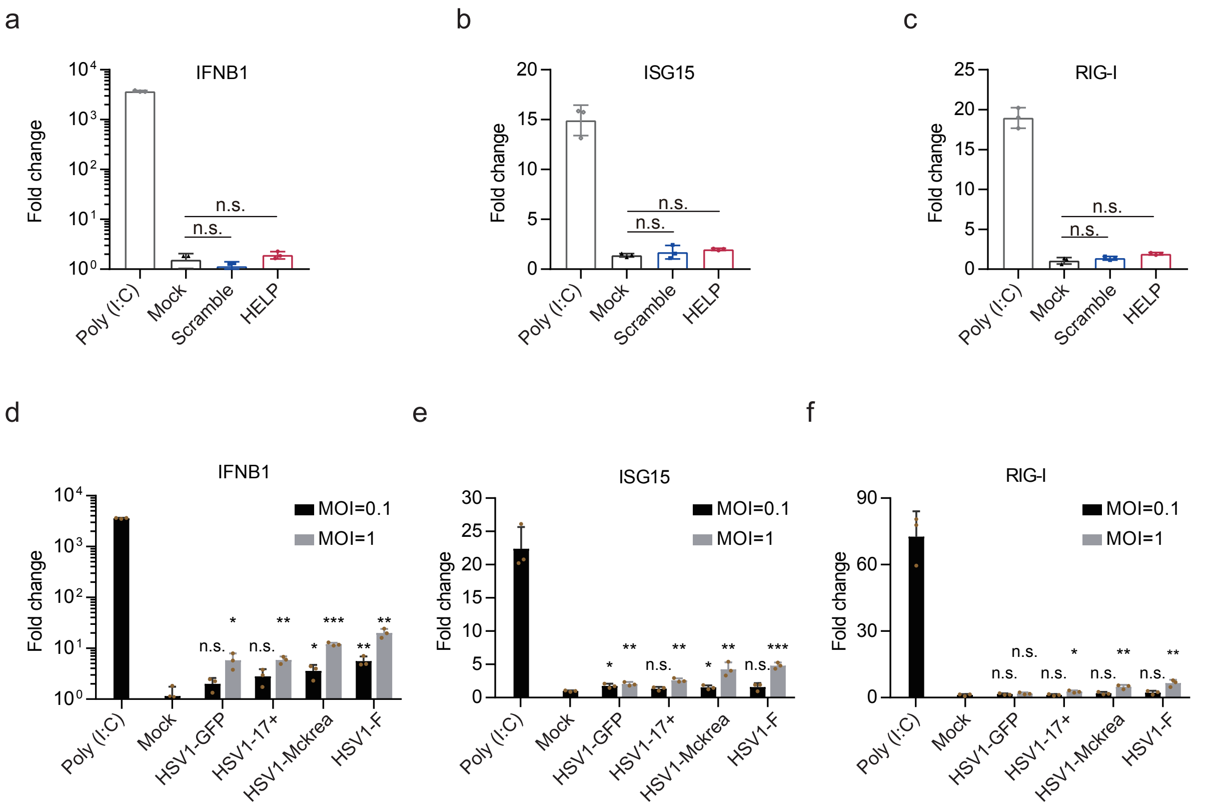


**Supplementary Figure 4. Innate immune response induced by HELP and different strains of HSV-1 in THP-1 derived macrophages.** Cells were harvested for IFNB1, ISG15 and RIG-I analysis by RT-qPCR 6 hr after transduction (**a-c**) or infection (**d-f**). Error bars represent ±s.e.m. Unpaired two-tailed Student's t-tests were performed, *<0.05, ***P*< 0.01, ****P*< 0.001, n.s.=non-significant.

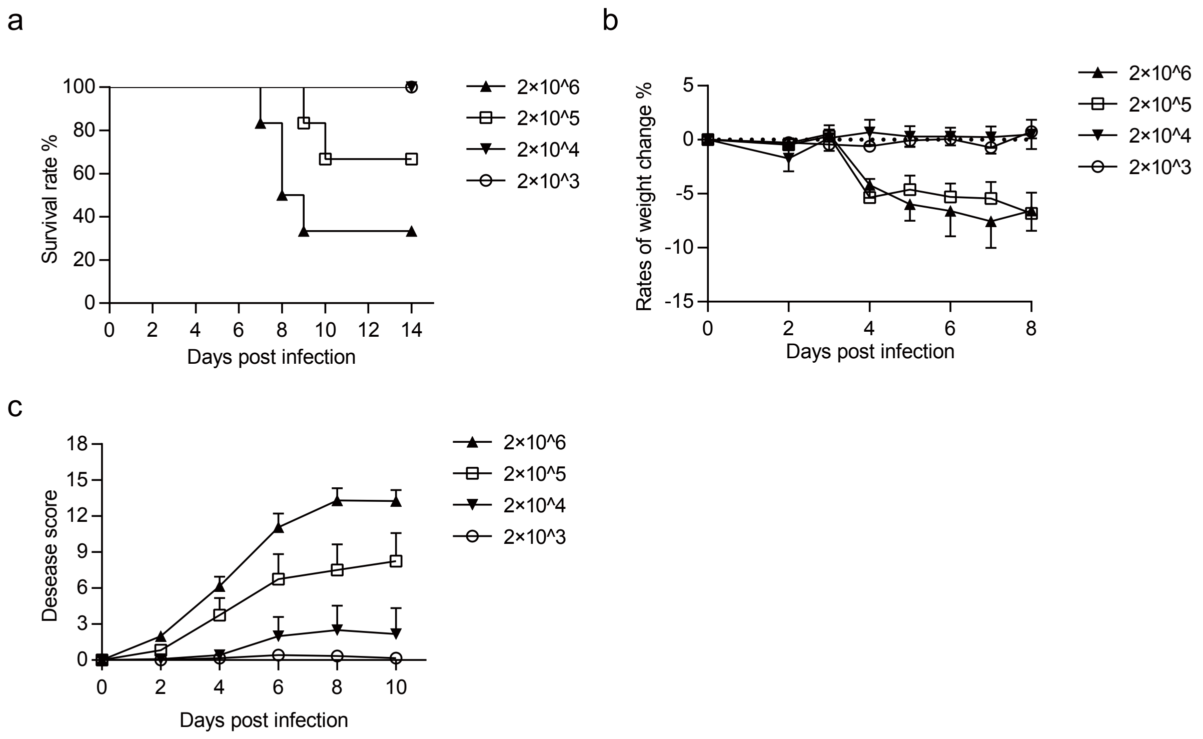


**Supplementary Figure 5. Dose-response and keratitis symptoms.** The mice were inoculated with different dosages of HSV-1 17syn+ on scarified corneas and recorded for survival rates, body weights and disease scores on the indicated days after infection.

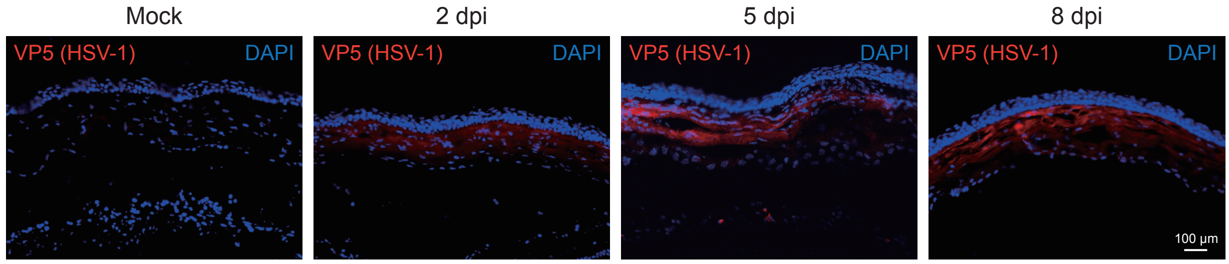


**Supplementary Figure 6. Time course of HSV-1 infection in the corneas.** The mice were inoculated with $2\times10^6$ PFU HSV-1 17syn+ on scarified corneas. Sections were prepared on 2, 5 and 8 days post-infection, respectively. dpi, day post infection.

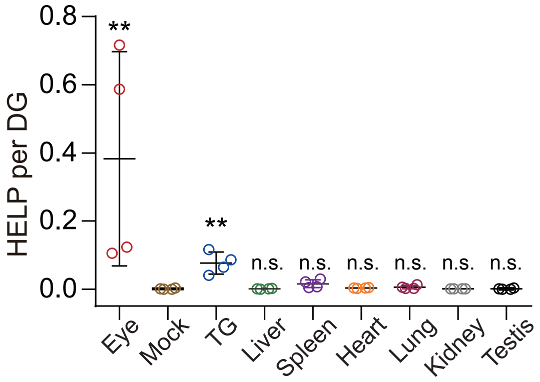


**Supplementary Figure 7. Tissue distribution of HELP in the whole body.** qPCR quantification of HELP dissemination in different tissues as viral genome (VG) per diploid genome (DG) (n=4 mice).

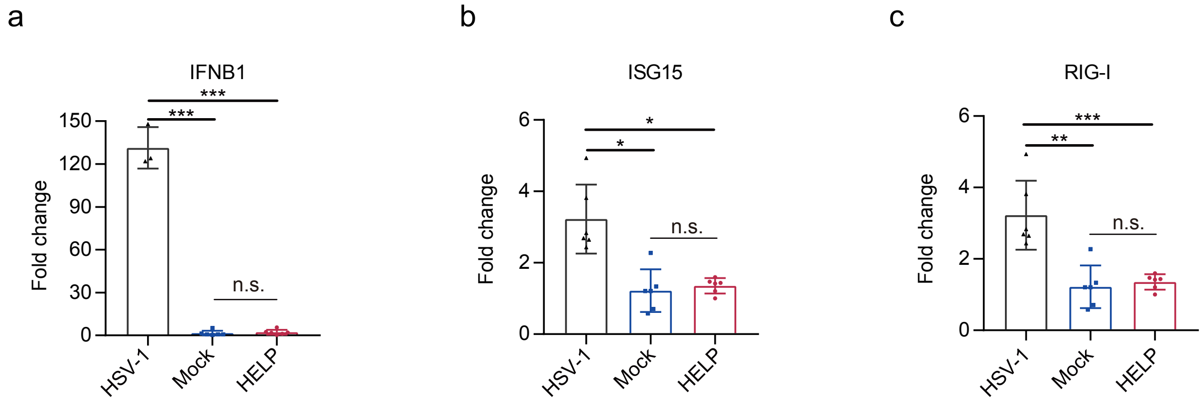


**Supplementary Figure 8. Innate immune response induced HELP and HSV-1 infection *in vivo*.** Corneas were harvested for IFNB1, ISG15 and RIG-I analysis by RT-qPCR 24 hr after HSV-1 17syn+ infection in corneas or intracorneal injection of HELP (n=5 mice). Error bars represent ±s.e.m. Unpaired two-tailed Student's t-tests were performed, n.s.=non-significant.

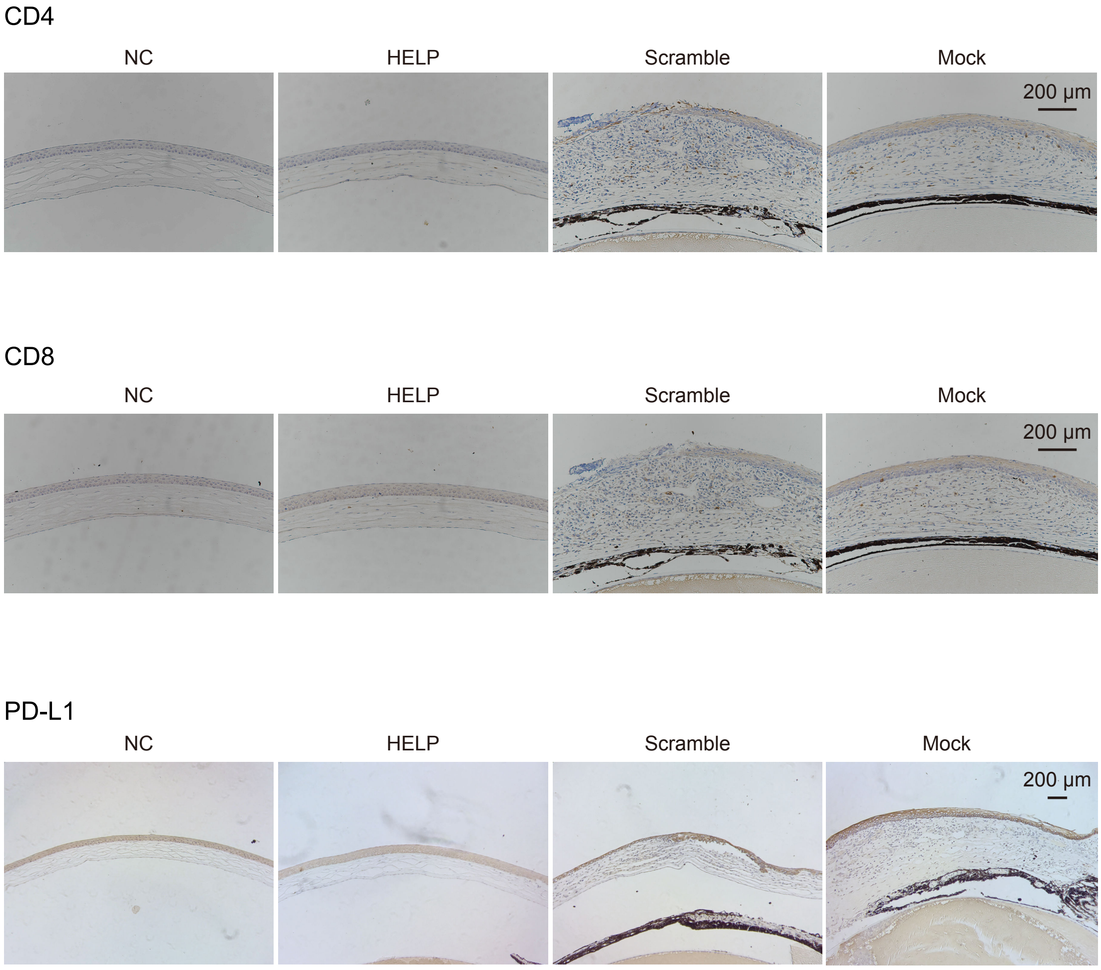


**Supplementary Figure 9. T cell infiltration and PD-L1 expression in the corneas *in vivo*.** Immunohistochemistry analysis of CD4+ and CD8+ cells infiltration and expression of PD-L1 in the corneas 14 days after infection.

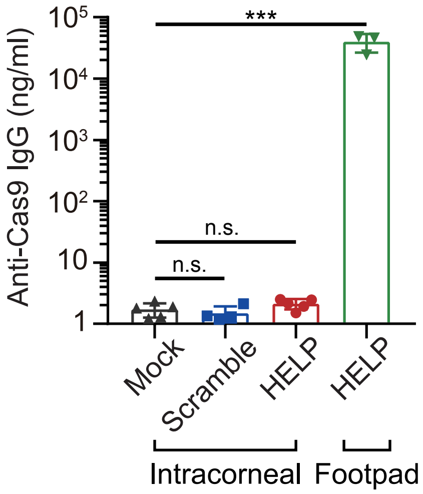


**Supplementary Figure 10. Cas9-specific IgG in the serum.** Mouse serum were collected at 14 dpi and analysed for anti-Cas9 IgG induction by HELP and non-targeting (scramble) mLP (n=5 mice). 200 ng p24 HELP and scramble or 2 μL PBS were injected into mice cornea by intrastromal injection. Footpad injection of 200 ng p24 HELP used as a positive control. Error bars represent ±s.e.m. Unpaired two-tailed Student's t-tests were performed, ***, $P<0.001$, n.s.=non-significant.
